# Extraction-free COVID-19 (SARS-CoV-2) diagnosis by RT-PCR to increase capacity for national testing programmes during a pandemic

**DOI:** 10.1101/2020.04.06.028316

**Authors:** Paul R Grant, Melanie A Turner, Gee Yen Shin, Eleni Nastouli, Lisa J Levett

## Abstract

Severe Acute Respiratory Syndrome coronavirus 2 (SARS-CoV-2) causes Coronavirus disease 2019 (COVID-19), a respiratory tract infection. The standard molecular diagnostic test is a multistep process involving viral RNA extraction and real-time quantitative reverse transcriptase PCR (qRT-PCR). Laboratories across the globe face constraints on equipment and reagents during the COVID-19 pandemic. We have developed a simplified qRT-PCR assay that removes the need for an RNA extraction process and can be run on a real-time thermal cycler. The assay uses custom primers and probes, and maintains diagnostic sensitivity within 98.0% compared to the assay run on a high-throughput, random-access automated platform, the Panther Fusion (Hologic). This assay can be used to increase capacity for COVID-19 testing for national programmes worldwide.

## Introduction

Coronavirus disease 2019 (COVID-19) is a respiratory tract infection caused by a newly emergent coronavirus – Severe Acute Respiratory Syndrome coronavirus 2 (SARS-CoV-2) – which was first recognised in Wuhan, Hubei Province, China, in December 2019. Genetic sequencing of the virus suggests that SARS-CoV-2 is a betacoronavirus closely linked to SARS coronavirus 1 (Wu et al.2020).

The standard molecular diagnostic test for this virus is a multistep process involving viral RNA extraction and real-time quantitative reverse transcriptase PCR (qRT-PCR). Although many companies have produced PCR kits to amplify the viral RNA, RNA extraction at any scale in a diagnostic laboratory is performed on a limited number of automated platforms that require specific reagents and consumables. This has led to significant effort to build large laboratories with existing research equipment to increase testing capacity, and to extract RNA on more open platforms that enable non-specific reagents and plastics to be used.

The COVID-19 pandemic placed severe constraints on the availability of laboratory equipment, reagents and consumables required for molecular diagnostics in the UK and Europe. This delayed the ability to scale-up testing capacity required for healthcare and population screening.

At Health Services Laboratories (HSL), we developed a qRT-PCR that can be run on a high-throughput, random-access automated platform, the Panther Fusion (Hologic). Using the Open Access facility on this platform, custom primers and probes designed in-house can be added to a DNA/RNA extraction cartridge. In London, this qRT-PCR was used for large-scale testing of patients hospitalized with suspected COVID-19. However, the COVID-19 pandemic also led to these cartridges being in short supply.

Using the same primers and probes, we have now developed a qRT-PCR that can be run on a real-time thermal cycler without the need for an RNA extraction process. This qRT-PCR maintains sensitivity to within 98.0% of the assay run on the Panther Fusion.

## Method

A panel of SARS-CoV-2 positive and negative samples was used to compare the RNA extraction and RNA-extraction free methods.

### RNA extraction

100 μL viral transport medium (VTM) from a swab was added to 100 μL Qiagen Lysis buffer containing guanidinum to inactivate the virus. This was then processed either on a QIAsymphony SP using the QIAsymphony DSP Virus/pathogen Mini kit and Complex 200 protocol, or the EZ1 Advanced XL using the EZ1 DSP virus kit. The elution volume was set to 60 μL, and 10 μL of the purified RNA was added to the PCR.

### Direct sample transfer

10 μL, 5 μL and 2 μL of sample expressed in viral transport medium was added directly to the PCR without any heating step, the plate was sealed and thermal cycling begun.

### Sample heating prior to direct transfer

50 μL of sample expressed in viral transport medium was added to a PCR tube and heated to 95 °C for 10 mins prior to loading into the PCR at 10ul, 5 μL and 2 μL.

### RT-PCR

A 20 μL reaction containing 10 μL RNA, 5 μL 4x TaqMan Fast Virus 1-step Master Mix (Applied Biosystems) and 5 μL primer and probe mix as shown in Table 1. Where VTM was added directly to the PCR at 5 μL or 2 μL, this was made up to 10 μL with RNase-Free water.

Cycling was performed at 56 °C for 15 min for reverse transcription, followed by 95 °C for 20 sec and then 45 cycles of 95 °C for 3 s, 60 °C for 30s using an Applied Biosystems QuantStudio 5 real-time PCR system (ThermoFisher Scientific).

**Table 1:**
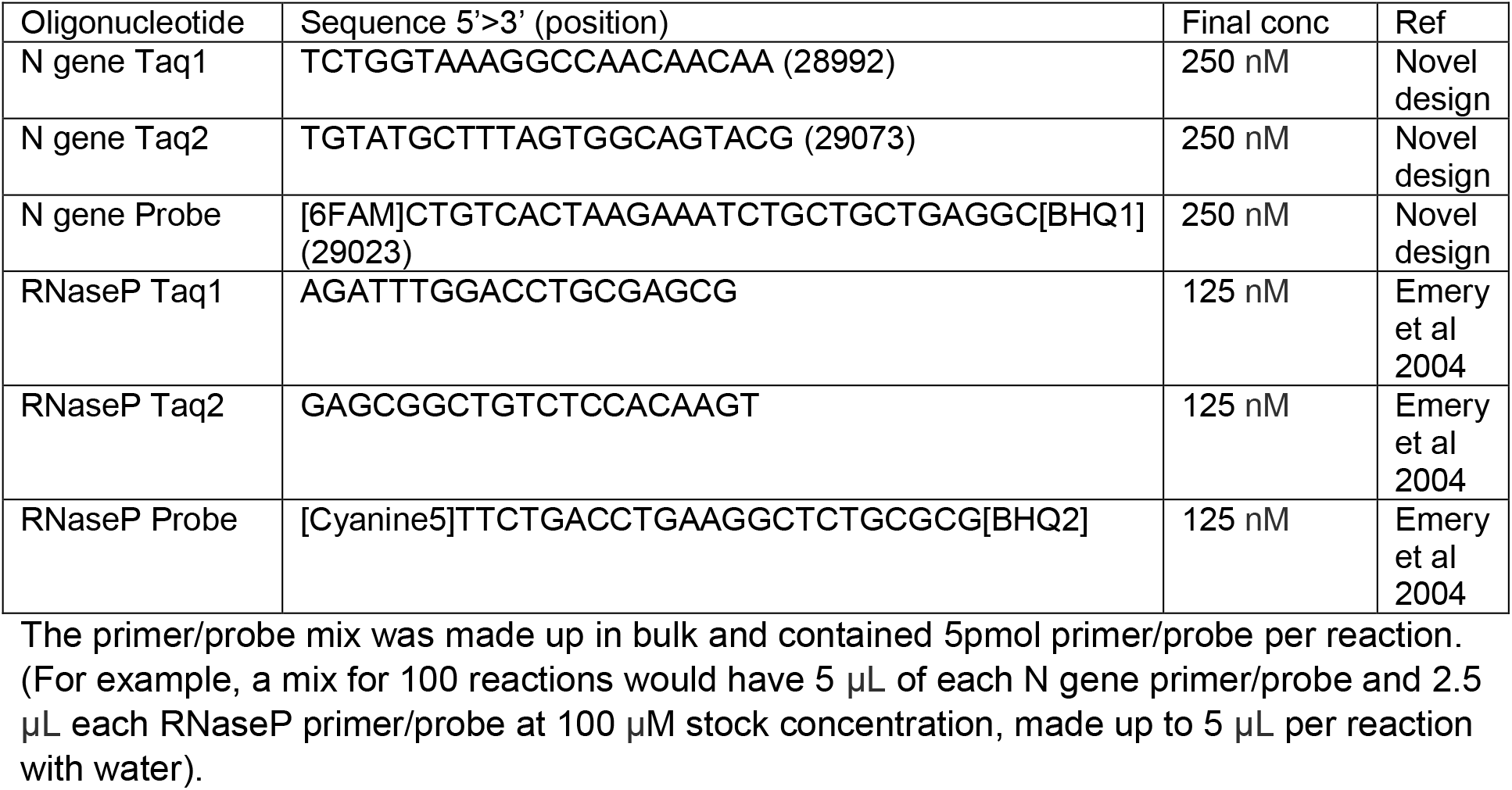
Primer/Probe sequences and concentrations

### Panther Fusion Open Access Assay

Panther Fusion PPR was made with the SARS-CoV-2 N gene primers and probe (Table 1) and Hologic internal RNA IC control primers and probe. A PPR tube for 40 reactions was made by adding 3 μL each N gene primer or probe (at 100 μM stock) and 8 μL of the RNA IC primer and probe (Supplied by Hologic). The salts were KCl at 100 mM (37.5 μL of 2M stock), MgCl_2_ at 3mM (4.5 μL of 1M stock) and Tris (pH 8.0) at 8mM (12 μL of 1M stock). This primer probe salt mix was made up to 1200 μL with water and overlaid with 400 μL mineral oil (Hologic) in a 2ml PPR tube (Hologic). This assay was run on the Panther Fusion (Hologic) with and Open Access RNA/DNA enzyme cartridge.

## Results

The RNA extraction method was compared to the direct addition of samples to the RT-PCR with and without prior heating.

When 10 μL of the heated or unheated sample was added to the PCR, no amplification was observed. Both the direct addition methods gave lower median Ct values than those added after heat treatment and were equivalent to the EZ1 (Qiagen) extraction (Figure 1).

**Figure 1:**
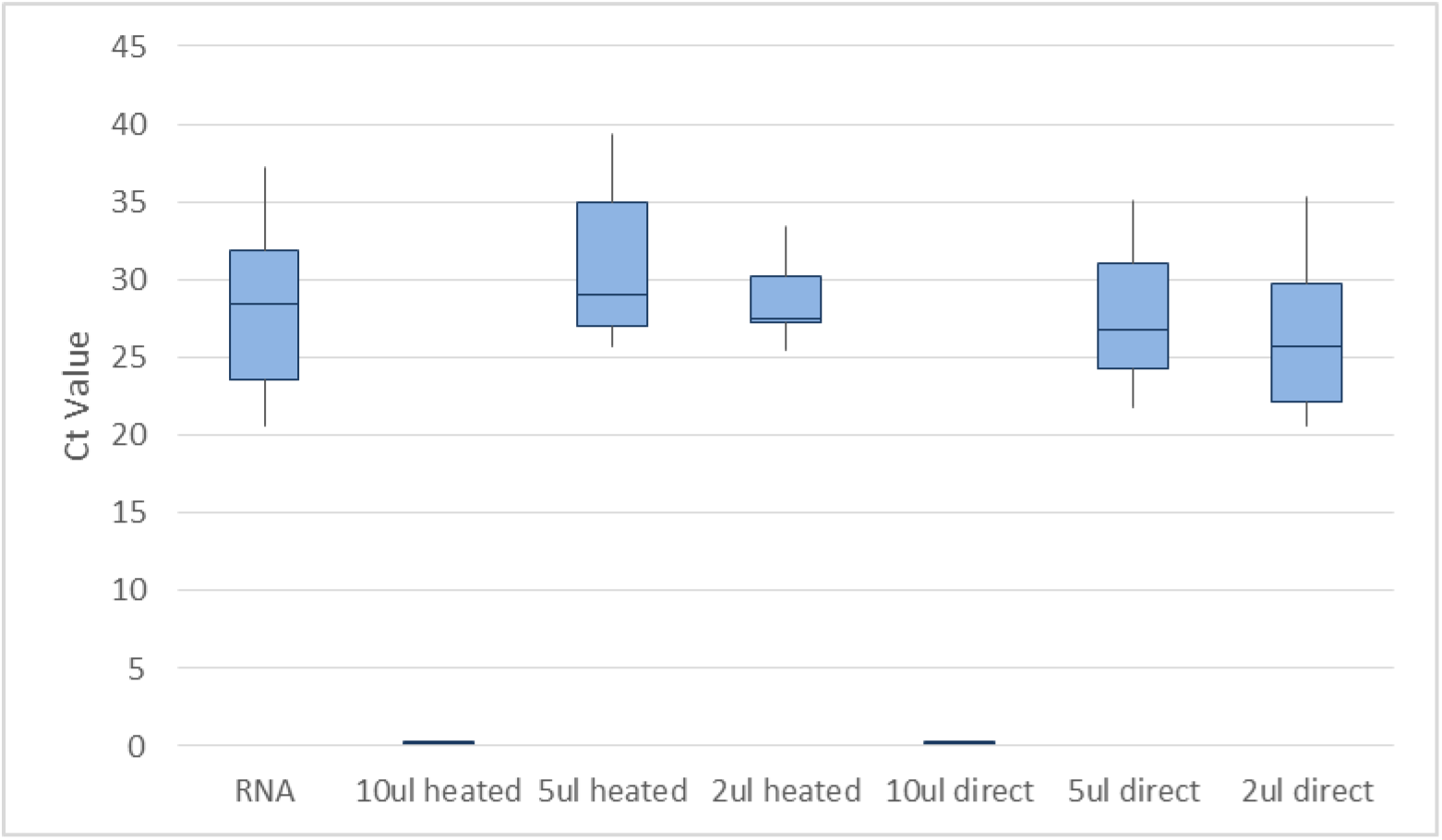
Median and interquartile range for Ct values obtained using different methods

The lowest Ct values were achieved by adding 2 μL of the VTM direct to the PCR without any prior heating (median Ct value 25.74 vs 29.51 using EZ1 RNA). This method was selected for further analysis.

The direct addition of 2 μL sample to the PCR was compared to the standard method in use within the clinical laboratory using the Open access channel of the Panther Fusion. An overall accuracy of 98.8% was achieved compared to the Panther Fusion assay (see Table 2).

**Table 2:** Diagnostic accuracy

|  | Results (n) |  |  |  | Test Performance (%) |  |  |  |  |
| --- | --- | --- | --- | --- | --- | --- | --- | --- | --- |
|  | TP | TN | FP | FN | sensitivity | specificity | PPV | NPV | Accuracy |
| 2 μL Direct | 96 | 71 | 0 | 2 | 98.0 | 100 | 100 | 97.3 | 98.8 |

TP = True Positive, TN = True Negative, FP = False Positive, FN = False Negative, PPV = Positive Predictive Value, NPV = Negative Predictive Value

The analytical sensitivity was compared by diluting a positive clinical sample to end point, and testing using the extracted RNA and adding VTM directly to the PCR. The results are shown in Table 3.

**Table 3:**
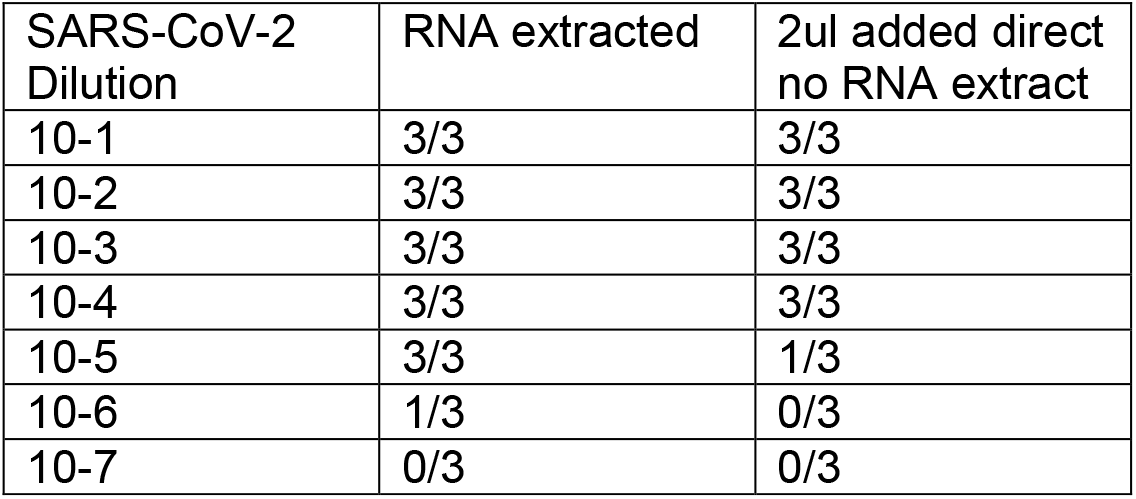
Analytical sensitivity

There was no cross reactivity with conventional coronavirus types OC43, NL63, 229E, HKU1, and a number of other respiratory viruses such as influenza A and B, respiratory syncytial virus, parainfluenza, metapneumovirus, adenovirus, rhinovirus and enterovirus.

## Conclusion

Direct addition of samples to the qRT-PCR without extraction with a diagnostic sensitivity of 98.0%, specificity of 100% and accuracy of 98.8% compared to the method on the Panther Fusion. This simplifies the process for COVID-19 testing, and will enable increased capacity in diagnostic laboratories.

## Discussion

Implementation of this method will enable laboratories to provide a COVID-19 testing service without the need for RNA extraction equipment, reagents and consumables. Turn-around times are similar to those of high-throughput RNA-based assays, and faster than a two-step RNA extraction and qRT-PCR. Capacity can be significantly increased without the extraction step but is dependent on the number of safety cabinets for swab processing and number of real time PCR thermal cyclers.

Heating at 56°C for 15 minutes causes SARS CoV (SARS coronavirus) to lose Infectivity (WHO, 2003). Health and safety assessments have been completed and the process has been deemed safe to perform with relevant precautions and safety practices. Samples can be processed in batches of 96, each batch takes 56 minutes to run the rtPCR on the thermal cycler, the rate limiting step being the swab processing. Lower numbers would be processed more rapidly, within an equivalent time to a point-of-care test. Standard swab processing can be automated to speed up the initial process on a large scale.

Many laboratories use real-time thermal cyclers, so this method can be used to increase national screening capacity without the need for other specialized equipment or RNA extraction reagents.

Applying an extraction-free PCR protocol as described here would avoid limitations on COVID-19 screening capacity in the UK and elsewhere caused by global PCR reagent supply constraints. We recommend this method is explored further by other medical laboratories using alternative PCR reagents to improve the resilience and capacity of virology laboratories during the pandemic. The sensitivity of the assay will be dependent upon the PCR efficiency, and so other PCR protocols will need to be carefully evaluated with this new approach.

